# Microbial populations hardly ever grow logistically and never sublinearly

**DOI:** 10.1101/2024.09.02.610774

**Authors:** José Camacho-Mateu, Aniello Lampo, Mario Castro, José A. Cuesta

## Abstract

We investigate the growth dynamics of microbial populations, challenging the conventional logistic model. By analyzing empirical data from various biomes, we demonstrate that microbial growth is better described by a generalized logistic model, the *θ*-logistic model. This accounts for different growth mechanisms and environmental fluctuations, leading to a generalized gamma distribution of abundance fluctuations. Our findings reveal that microbial growth is never sublinear, so they cannot endorse—at least in the microbial world—the recent proposal of this mechanism as a stability enhancer of highly diverse communities. These results have significant implications for understanding macroecological patterns and the stability of microbial ecosystems.

## I. INTRODUCTION

Ever since Verhulst proposed it in 1838 [1], the logistic growth equation has become a standard in modeling populations in ecology—so much so that it has become a common default assumption when devising any population framework. Its rationale is simple—the per capita growth is assumed to be proportional to the fraction of resources left by the rest of the population. However, in Murray’s words [2], the logistic equation “is more like a metaphor for a class of population models with density-dependent regulatory mechanisms … and must not be taken too literally”. In fact, it is not difficult to figure out mechanistic benchmarks leading to other types of density-dependent growth [3, 4]. Tumors, for instance, are better modeled with a Gompertz’s law than with a logistic model [5, 6].

Whether population growth is logistic or not may have important consequences for ecological communities. For instance, in a recent work, Hatton et al. [7] show that a sublinear growth (a growth scaling with biomass *x* as *x*^*µ*^, with 0 *< µ <* 1) might resolve the longstanding stability-diversity controversy [8, 9]. Even though this result has been contested [10], there is no doubt that the growth law may influence important aspects of ecological communities. Furthermore, empirical evidence accumulated over decades [11, 12] suggests that the growth laws governing populations of birds, mammals, fish, and insects are indeed heterogeneous.

The question becomes particularly relevant for microbial communities because simple mechanisms of cellular growth that consider both excluded volume and cell diffusion [3] generate effective growth laws other than the logistic model. Until the arrival of metagenomics, determining how microbial populations grow was out of the question. At present though, the unprecedented amount of available data [13–15] renders the problem amenable to analysis. Whether or not microbial populations grow logistically can be crucial to explain macroecological patterns observed in these communities [16–23] because some of them seem to depend critically on the logistic assumption [16].

In this work we will show (*i*) that the microbial growth in several real biomes [24] is better described by a generalization of the logistic model; (*ii*) that it does affect the emergent macroecological patterns; (*iii*) that, in spite of this, the proposed explanation for these patterns [16] can be ‘saved’ if properly stated; and (*iv*) that microbial growth is never sublinear—discarding any alleged benefit on the stability of microbial communities [7, 10].

## II. GENERALIZED GROWTH MODEL FOR MICROBES

A generalization of the logistic model that covers a wealth of different growth mechanisms is the *θ*-logistic model [25], defined by the equation

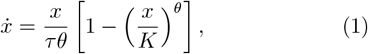

where *x* represents the population of a species (also referred to as abundance, or density of individuals), *τ* is its intrinsic growth time, and *K* is the carrying capacity. The parameter *θ* governs the dependence of the per capita growth rate with the abundance. The exponent *θ* encapsulates the effect of the different growth processes [3, 4]. A value *θ >* 0 describes a linear growth for small abundances 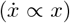—with *θ* = 1 representing the logistic model—whereas −1 *< θ <* 0 describes a sublinear growth 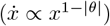. The transition from *θ >* 0 to *θ <* 0 is smooth because in the limit *θ* → 0 the equation recovers Gompertz’s model [26]

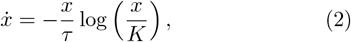

commonly employed to describe tumor growth [5, 6].

However, real biomes are subject to environmental changes. The rapid fluctuations they cause in species abundance calls for a stochastic modeling [16, 18, 28, 29]. This is achieved by adding a noise term to the *θ*-logistic model (1). Analysis of the abundance fluctuations in empirical data suggests introducing a multiplicative noise proportional to *x*, as in Grilli’s stochastic logistic model (SLM) [16, 18, 30, 31] (see [32], Fig. S1). The result is the stochastic *θ*-logistic model (S*θ*LM) [4], defined by the Langevin equation

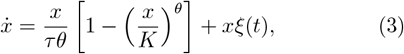

where *ξ*(*t*) is a zero-mean, Gaussian, white noise such that ⟨*ξ*(*t*)*ξ*(*t*^*′*^)⟩ = *w*δ(*t* − s*t*^*′*^), *w* being the noise variance. Figure 1b) illustrates typical realizations of the S*θ*LM for different values of *θ* (*θ* = 1 reproduces the SLM).

**FIG. 1.**
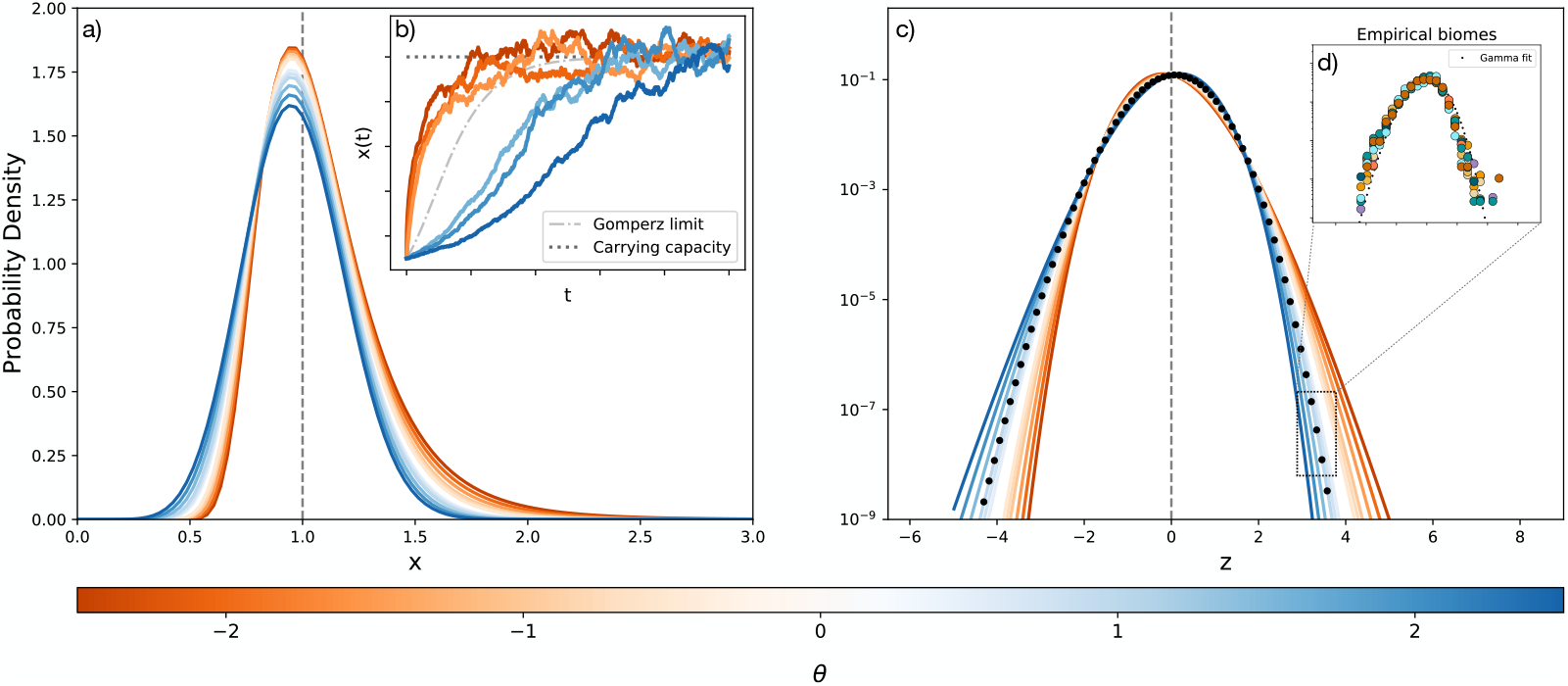
a) Generalized gamma corresponding to the abundance fluctuation distribution *P* (*x*), Eq. (7), for several values of *θ* (indicated by the colorbar). b) Realizations of the S*θ*LM (3) for *θ* = −2.5, −2, −1.5, 1.5, 2, 2.5, were obtained using an Euler-Maruyama integration scheme [27] with initial condition *x*(0) = 0.1 and integration time-step Δ*t* = 10^−3^. The black dotted line signals the value of the carrying capacity *K*, while the grey dot-dashed line provides the solution of Eq. (2) (corresponding to Gomperz’s limit *θ* → 0). c) Graphical representation of log 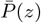 [c.f. Eq. (16)], i.e. the logarithm of the abundance fluctuation distribution expressed as a function of the standardized abundance *z* [c.f. Eq. (10)], for several values of *θ*. The black dots portray the fit to the biomes of inset d), where we illustrate log 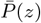 for nine real biomes: Seawater, Soil, Sludge, Oral1, Gut1, Gut2, Glacier, River, Lake [24]. All plots have been performed with *τ* = 0.1, *K* = 1, *w* = 0.5.

Equation (3) describes the intrinsic stochastic dynamics of the abundance of different microbial taxa, each of which is expected to be described by specific parameters. For the sake of clarity though, no taxonomic identity is explicitly included in the equation.

## III. ABUNDANCE FLUCTUATION DISTRIBUTION

The probability distribution of the S*θ*LM (3) in the steady state (the so-called abundance fluctuation distribution, AFD) can be readily obtained through its asso-ciated Fokker-Planck equation *P*_*t*_ + *J*_*x*_ = 0, where

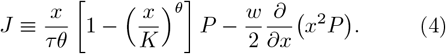

In the steady state, the current is constant. This constant is zero because of the boundary conditions *P* (0) = *P* (*x*→ ∞) = 0, so we can find the stationary probability *P* (*x*) as the solution of

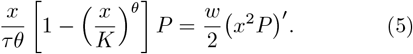

Denoting *q*(*x*) = *x*^2^*P* (*x*), this equation becomes

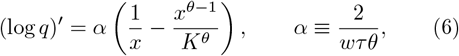

hence

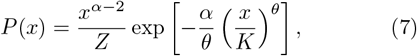

where

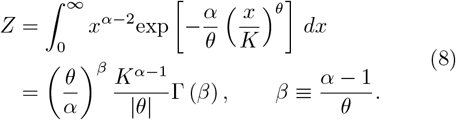

It is important to notice that *β >* 0 for any *θ <* 0, as well as for any 0 *< θ <* 2*/wτ*.

Probability density (7)—whose shape for different values of *θ* is illustrated in Fig. 1—describes how the abundance of a species fluctuates as a consequence of environmental noise. In the context of macroecology [33, 34] this is one of the patterns that characterize microbial communities. As a matter of fact, it has been recently proposed that this AFD has the shape of a gamma distribution [16], which corresponds to the case *θ* = 1 (logistic growth). Instead, when *θ ≠* 1, the AFD (7) has the shape of the generalized gamma distribution [4], whose moments are given by

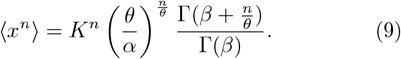

A generalized gamma is a versatile distribution. Apart from recovering the gamma distribution for *θ* = 1, it can also adopt other familiar shapes for different parameter choices. For instance, it can be a stretched exponential (*α* = 2), an ordinary exponential (*α* = 2, *θ* = 1), a (positive) half-normal (*α* = *θ* = 2), or a Weibull (*α* = *θ*). In the limit *θ* → 0 it becomes a lognormal (see Appendix A). Furthermore, for *θ <* 0 it abruptly drops to zero for small *x* whereas it decays as a power law for large *x*.

As the gamma shape of the AFD is an empirically well-established macroecological pattern in microbial communities [16, 21–23], the previous result seems to rule out any other growth mechanism but the logistic one. Before we jump to a wrong conclusion though, let us notice that all empirical analyses of the AFD of microbial species use the standardized variable

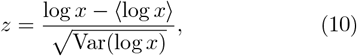

so it is 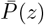, the probability density of this standard variable *z* that we need to use for comparison.

In order to obtain 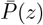 let us introduce the intermediate variable

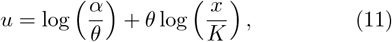

whose probability density relates to (7) through 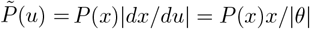. Thus,

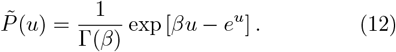

We can now relate *u* to *z* by noticing that

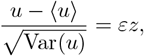

where *ε* sign(*θ*). In order to calculate *u* and Var(*u*) we use the identity

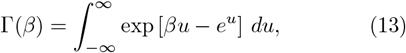

in terms of which

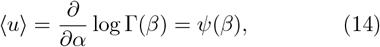

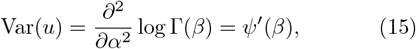

where *ψ*(*x*) denotes the digamma function [35]. Thus, as 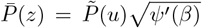, we conclude that 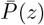 takes the form of an exp-gamma distribution, namely

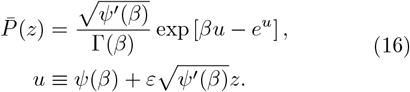

This distribution is depicted in Fig. 1c for different choices of *θ*. Figure 1d also shows good agreement with empirical AFDs.

Two remarkable features of distribution (16) are worth noticing: first of all, it is independent of *K*, and secondly, except for the sign of *θ* it depends on the rest of the parameters (*τw* and *θ*) only through the combination *β*. Hence, the value of *β* obtained by fitting 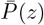 *s* to a set of data, can be interpreted in very different ways—either we assume *θ* = 1 and use *β* to determine *τw*, or we fix *τw* and use *β* to determine *θ*. In other words, the fact that (16) is a good description of empirical AFDs does *not* rule out a *θ*-logistic growth. Previous macroecological patterns are perfectly compatible with alternative growth laws.

We still have to discuss the dependence on *ε* = ±1. This parameter discriminates the two classes of growth at low abundances: linear (*ε* = +1) and sublinear (*ε* = −1). The skewness coefficient of 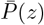 is (see Appendix B)

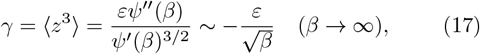

so −*ε* determines the sign of the skewness of this distribution. Notably, all empirical distributions are negatively skewed, hence supporting the value *ε* = +1, i.e. linear growth (see Fig. 1c). This rules out—for microbes—the sublinear growth proposed by Hatton et al. [7] as a stability enhancer of highly diverse communities.

Unfortunately, the same feature that makes of 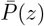 a universal macroecological pattern (the dependence through the single parameter *β*) renders it useless to discriminate the kind of growth law followed by the different microbial species. In order to achieve this goal, we need to fit directly Eq. (7) to the data of microbial abundances.

## VI. BAYESIAN INFERENCE OF THE GROWTH PARAMETER

Estimates of the parameter *θ* have been obtained in the past through least squares fits of the dynamic equation (1) to time series data [7, 12, 25, 36]. This method has many drawbacks—such as the poor estimation of derivatives produced by a finite-difference scheme applied to noisy data, or the underlying assumption that the noise follows a normal distribution. The estimates of *θ* are thus unreliable. Instead, we took advantage of the knowledge of *P* (*x*) [c.f. Eq. (7)] and performed a Bayesian inference of the parameter *θ* [37, 38].

Databases of microbiomes do not provide lists of species abundances. A typical sample is a vector of counts **n**_*s*_ = (*n*_1,*s*_, *n*_2,*s*_, …, *n*_*S,s*_), where *n*_*i,s*_ is the number of reads attributed to taxon *i* = 1, …, *S* in sample *s*. The total number of reads across all taxa in each sample, Σ_*i*_ *n*_*i,s*_ = *N*_*s*_, is referred to as ‘sampling effort’ or ‘sampling depth’. Thus, we must add the variability introduced by the sampling process to the inherent variability of the abundances induced by the dynamics [captured by the distribution (7)]. We can describe this sampling by a Poisson distribution with mean *x*_*i,s*_*N*_*s*_. Therefore, the probability to observe *n*_*i*_ counts of the *i*th taxon in the given sample *s* is obtained by marginalizing over *x*_*i*_, namely

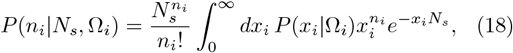

*P* (*x* |Ω) being the generalized gamma distribution (7)— which for the sake of simplicity we parametrize as in the R package dgen.gamma, i.e.

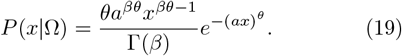

Here Ω denotes the set of parameters *β, a, θ*, where

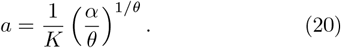

Now, the posterior distribution of parameters Ω_*i*_ can be obtained as

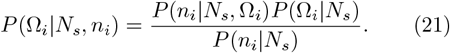

Guided by the maximum entropy principle [37], we use exponential distributions as priors for the parameters:

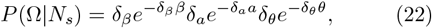

with δ_*β*_, δ_*a*_, δ_*θ*_ *>* 0 small constants. These priors ensure the positiveness of these parameters but are otherwise quite agnostic with respect to what their values are.

The method just described produces an estimate of the value of *θ* for individual species. However, a comparison of the estimates for different species suggested that a gamma distribution

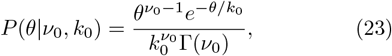

for an appropriate choice of the “hyper-parameters” *ν*_0_, *k*_0_, might be a reasonable model for the distribution of *θ* across species. Accordingly, to estimate this distribution more accurately we adopted a global strategy, inferring the parameters for all species at once, using a hierarchical Bayesian model. In other words, we calculate the global posterior

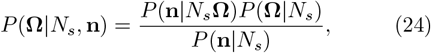

where **n** is the vector of counts of all species, and **Ω** denotes the set of parameters of all species plus the hyperparameters *ν*_0_, *k*_0_. Then we choose the prior

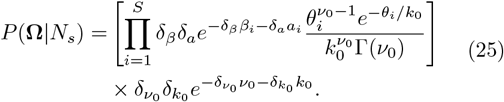

where we have also used exponential priors for the hyperparameters.

This hierarchical Bayesian approach not only refines the estimates but also allows cross-learning among taxa, enhancing the overall robustness and accuracy of our parameter estimations while reducing overfitting at the same time. Implementation details can be found in Appendix C.

An important issue related to Bayesian inference is identifiability. Roughly speaking, we say that a parameter is identifiable if its posterior distribution is relatively narrow and peaked. We have conducted a synthetic test to prove that the parameters of the S*θ*LM can actually be inferred by the described procedure. Thus, we generated a random sample of size *N*_*c*_ and fixed sampling depth *N*_*s*_ from a synthetic population consisting of *S* = 4 independent taxa. We then applied the Bayesian inference method just described, assuming that abundances follow the generalized gamma distribution (19) with equal parameters *β* = 2 and *a* = 0.2, but different *θ* parameters (*θ* = 0.5, 1, 1.5, 2). Given that there are only four taxa, the *θ*s are inferred individually—using a prior 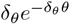. The high reliability in recovering these parameters exhibited by the method (as illustrated in Figure 2) is reassuring.

**FIG. 2.**
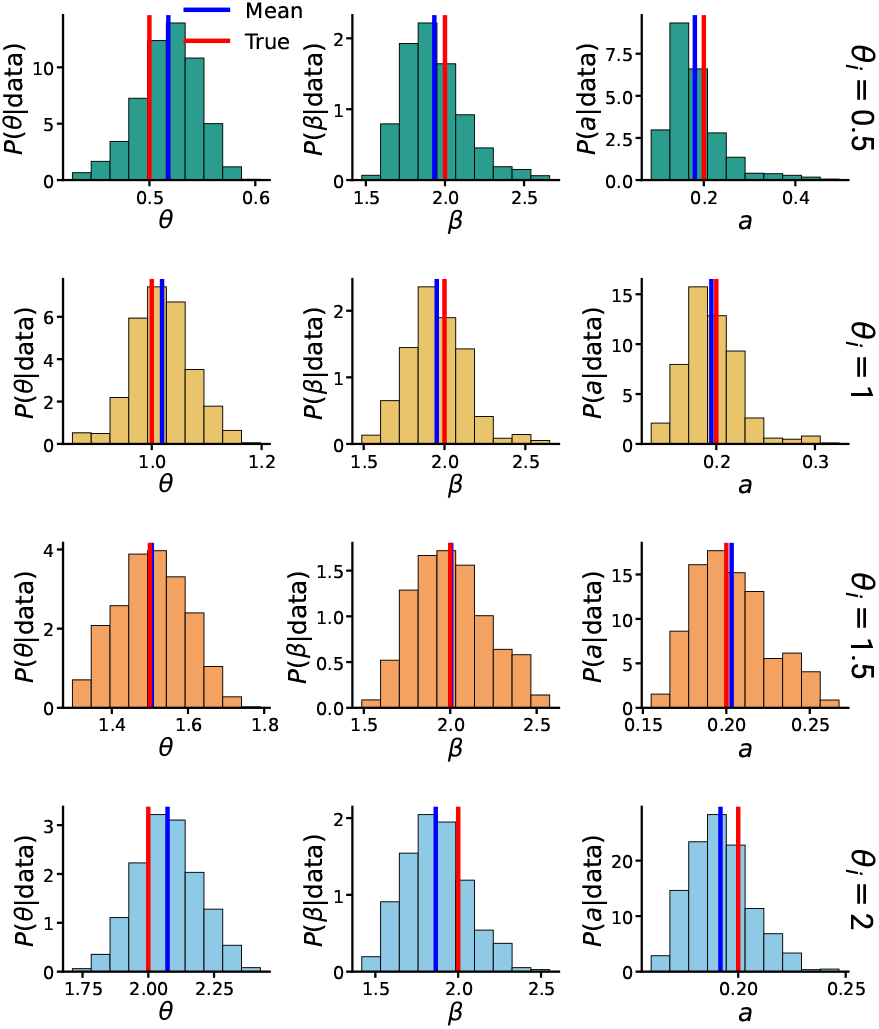
Marginal posterior distributions for parameters *θ, β*, and *a* [see Eq. (19)] as obtained for a synthetic data set consisting of *S* = 4 taxa with *θ* = 0.5, 1, 1.5, 2, and the same values of *β* = 2 and *a* = 0.2. The true parameter values are marked by red lines; the mean of the posterior predictive distributions [Eq. (26)] by blue lines. The narrow and peaked posteriors around the true values prove the identifiability of these parameters using Bayesian inference. Total number of samples *N*_*c*_ = 15000, sampling depth *N*_*s*_ = 5000.

Figure 3 shows the distributions for *θ* for several biomes estimated as we have just explained. The gamma functions plotted in this figure correspond to the inferred maximum posterior hyperparameters. For comparison, the value *θ* = 1 (logistic growth) is marked with a dashed vertical line. The figure reveals that logistic growth is hardly the most probable growth law among microbes— the values of *θ* ranging from nearly zero to *θ* ≈5.

**FIG. 3.**
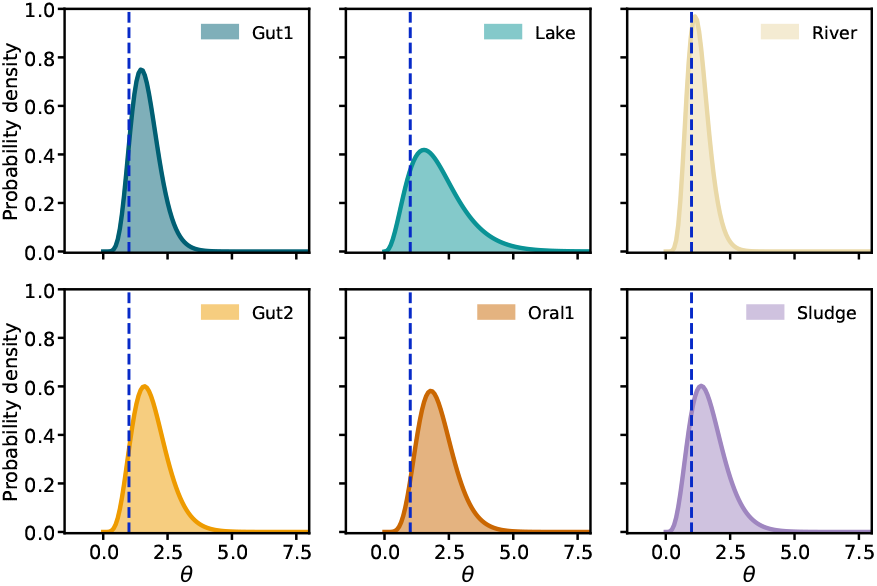
Gamma distributions of *θ* across species for several biomes as obtained from the estimated maximum posterior hyperparameters. Dashed blue lines mark the value *θ* = 1 corresponding to logistic growth. For computational limitations, in making this figure we only included species appearing in more than 95% of the samples.

To assess the quality of our Bayesian inference, we checked that the outcome of the posterior predictive agrees well with the AFD of individual species (see [32], Figs. S6–S7). Posteriors predictive are obtained as

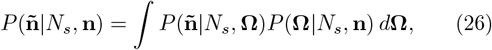

where the integral covers the whole parameter space.

These distributions can be sampled by Monte Carlo simulations if we interpret the integral above as the average of the likelihood *P* (**ñ**|*N*_*s*_, **Ω**) over the posterior distribution *P* (**Ω** |*N*_*s*_, **n**).

We can illustrate the accuracy of this posterior predictive by comparing averages (Fig. 4a) and variances (Fig. 4b), as obtained from (26) and from empirical data. As a reference, the insets in these figures provide the results obtained assuming that the AFD follows a gamma distribution (as derived from the logistic, *θ* = 1, growth).

**FIG. 4.**
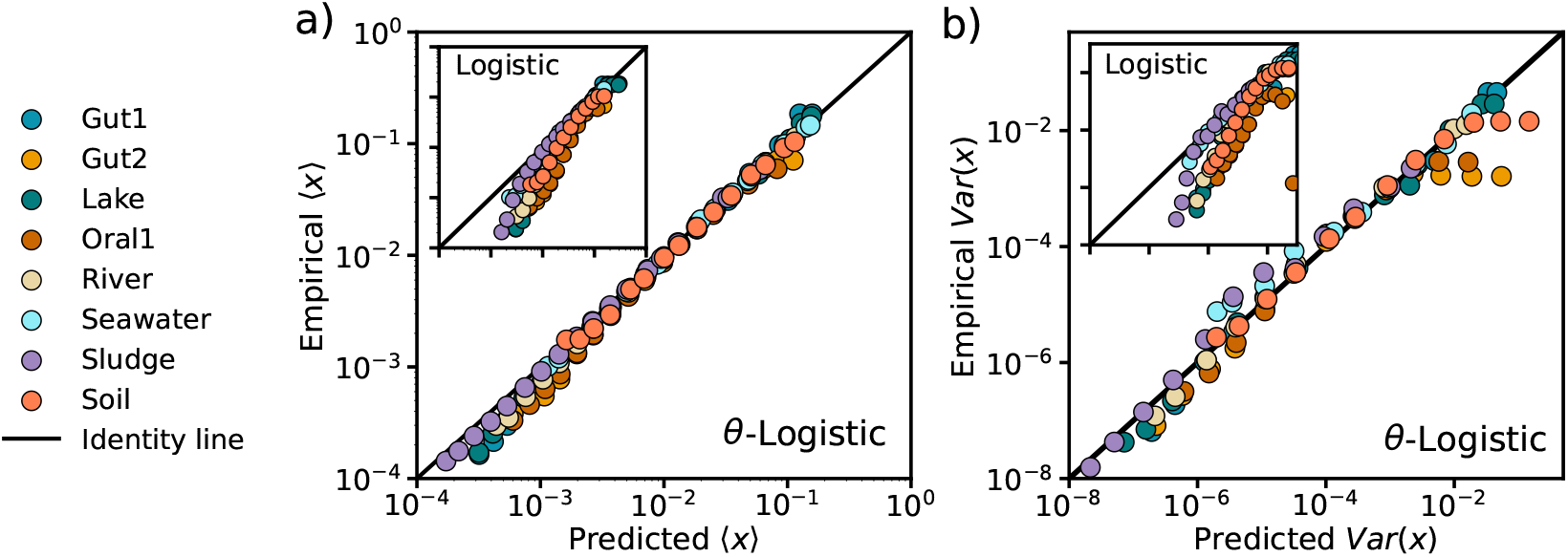
Plot of a) the abundance average; and b) the variance for different biomes [24], as obtained from Eq. (9) with the inferred parameters, vs. their corresponding empirical values. Dots provide a coarse-grained visualization of these magnitudes, with error bars intentionally left aside for the sake of clarity (see [32], Figs. S2–S5). Insets show the same test applied to the SLM (with logistic growth). Its tendency to underestimate both magnitudes across most of the population enforces the conclusion that microbes rarely grow logistically.

As a final test, we computed the distribution of the logarithmic fold-change *λ* = log(*x*_*s*+1_*/x*_*s*_) of two consecutive samples (*x*_*s*_ and *x*_*s*+1_) of the abundance of a given species. This distribution has been proposed as another macroecological pattern and has been the focus of interest of several recent studies [17, 20, 39, 40]. It is straightforward to derive it from the AFD provided the two samples are sufficiently uncorrelated. The result is (see Appendix D)

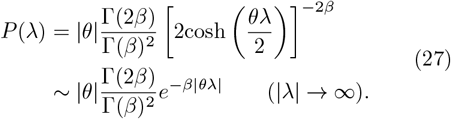

Interestingly, this is the distribution reported for *λ* when it is averaged across all samples and taxons [17]. In Fig. 5 we compare *P* (*λ*) as obtained from the inferred parameters with the empirical one. For each taxon, we calculate the maximum a posteriori estimate (the mode of the posterior distributions) and the 95% confidence interval. A more thorough comparison can be found in [32], Figs. S8–S9.

**FIG. 5.**
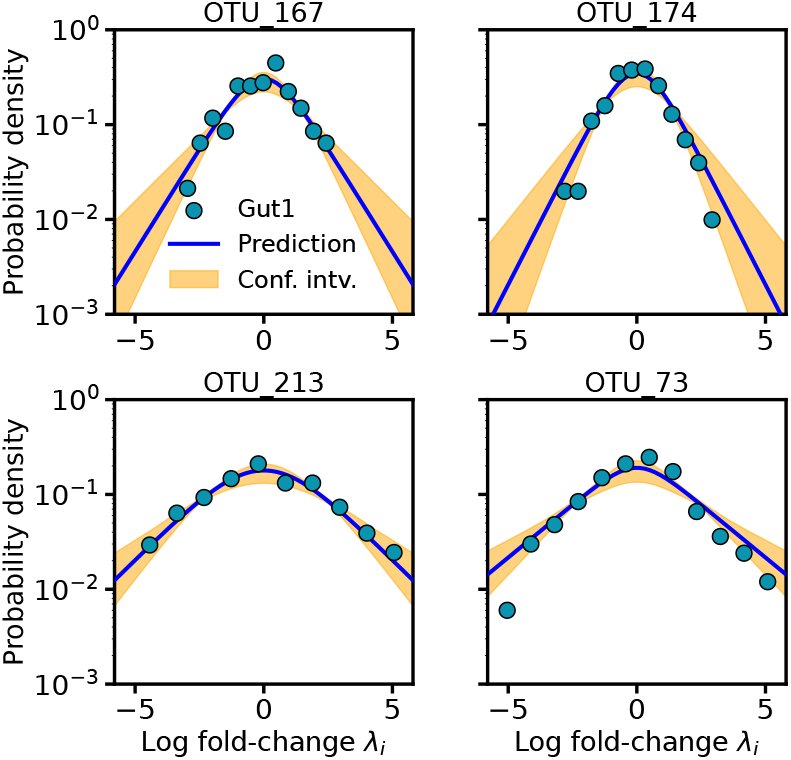
Logarithmic fold-change (see main text) distribution (27) for four randomly chosen species of the Gut1 microbiome [24]. Lines are the inferred distributions; orange regions span 95% confidence intervals for each *λ* value; dots are otained from empirical data. The good agreement assesses the accuracy of the AFD in reproducing the data. (In cross-sectional data, where autocorrelations are negligible, this distribution follows from the AFD; see Appendix D.)

## V. DISCUSSION

We conclude from our previous analyses that there is sufficient evidence to rule out logistic growth among microbes in favor of more sophisticated mechanisms, in line with previous evidence obtained from birds, mammals, fish, and insects [11, 12]. Deviations from the logistic growth can originate in excluded volume effects [3], collective behavior (cooperation or competition) of bacteria [4], difficult access to external resources of microbes living at the interior of a colony, or any other complex dynamics. This fact brings about a crucial change in an important macroecological pattern: the AFD takes the form of a generalized gamma distribution instead of the gamma shape that had been claimed so far [16, 21–23]. This does not mean that previous analyses were incorrect, because all of them were performed using the variable *z* [c.f. Eq. (10)]. As we have shown in this paper, in terms of this variable all generalized gammas fall into two distinct patterns—that are otherwise indistinguishable. This is the reason why all previous research has misinterpreted the shape of the AFD as a gamma distribution. Interestingly, all microbes seem to fall into just one of the two patterns. This rules out the possibility of sublinear growth, and therefore of a possible mechanism to enhance the stability of highly diverse ecosystems [7]. Thus, whether this is a plausible mechanism or just an artifact as it has been recently argued [10], is a controversy that has no relevance for microbial communities.

Our findings impact another common macroecological pattern: Taylor’s law. It has been argued [16] that the variance and the mean across microbial species are related by the power law 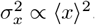. In a previous work [22] we showed that interactions introduce significant variability, albeit this law represents a general trend. Different growing mechanisms introduce an extra source of variability. As a matter of fact, given the expression (9) for the moments of the AFD, the ratio 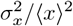 depends on *θ*, and this contributes an additional source of variability to Taylor’s law. How much of the variability observed in empirical data is due to this effect or to the existence of interactions is a question that deserves to be explored.

We have performed our analysis on the available microbial datasets as if species were isolated from each other. We are aware that the presence of interactions might impact the growing behavior of each species and provide an alternative explanation for the non-logistic growth and it is not unreasonable to argue that the parameter *θ* could be an effective proxy for those interactions. Addressing this question theoretically is problematic because inferring interactions is already a complex problem by itself [23, 41, 42]. Accordingly, the best approach to decide on the growth behavior of microbes would be to carry out specific experiments to determine how populations of different microbial species grow in isolation from the rest of the species—as opposed to how they do within their communities.

We cannot resist adding a few thoughts on the problem of the stability of highly diverse microbial communities. Having ruled out the alleged mechanism based on a sublinear growth, and taking into account different alternatives proposed in the literature [43– as well as recent theoretical speculations on microbial ecosystems [49], we advocate for the spatial distribution as the most plausible mechanism favoring the increase of diversity observed in microbiomes. A spatial distribution of microbes can be regarded as a metacommunity, where many smaller communities coexist. Their small size reduces competition, and the distribution of species across them enhances biodiversity [48, 49]. Whether this can be proved to be a valid mechanism, theoretically and empirically, is a very interesting open problem.

We would like to finish with a word of caution. In line with Murray’s remark on the logistic equation, and despite the analyses we have presented here, we do not think that Eq. (3) should be taken too literally either. The equation is nothing but an effective description of microbial dynamics that is more accurate than the simple logistic law. This means that we should not worry too much about attributing a biological meaning to the parameter *θ*. Microbial models are no more—but not less—than useful tools to describe real ecosystems. Nonetheless, they provide valuable insights about bacteria interactions and spatial organization, so we hope that this work helps trigger new experiments beyond the recollection of species abundances.

## Supporting information

Supplementary information of the text

## ACKNOWLEDGMENTS

We sincerely thank Dr. Pablo Catalán for very fruitful discussions, and for his help with the related bibliographic research on the biological significance of the *θ* parameter and the microscopic mechanisms behind population growth. This work has been supported by grants PID2022-141802NB-I00 (BA-SIC), PID2021-128966NB-I00, and PID2022-140217NB-I00, funded by MICIN/AEI/10.13039/501100011033 and by “ERDF/EU A way of making Europe”.

## Appendix A: The Gompertz limit

We shall derive this limit from Eq. (16). When *θ* → 0 we have *β*→ ∞. In this limit we can use the asymptotic expansions

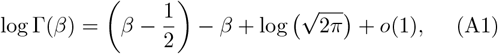

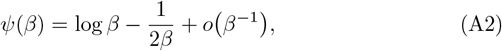

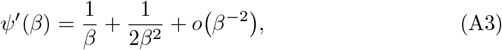

from which *u* and *e*^*u*^ become

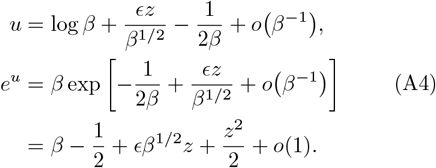

Therefore

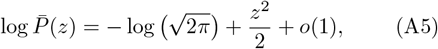

the equation of a lognormal distribution.

## Appendix B: Skewness of the exp-gamma distribution

Given that ⟨*z*⟩ = 0 and Var(*z*) = 1, the skewness coefficient of 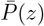 can be obtained as

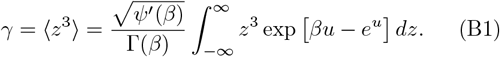

In terms of the variable 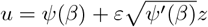

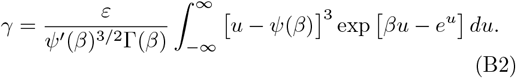

Finally, since [cf. Eq. (13)]

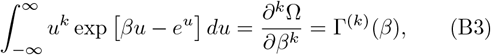

then

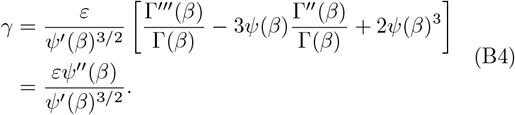

## Appendix C: Bayesian inference implementation

Bayesian inference was performed in *R* using *JAGS*, a software for analysis of Bayesian graphical models using Gibbs sampling [38]. *JAGS* is a very intuitive software that allows chaining distributions of different parameters that depend on each other, so it does not require us to know a closed expression for the likelihood (18). Figure 6 is a pseudo-code of the *JAGS* algorithm employed in the hierarchical Bayesian inference described in Sec. IV.

**FIG. 6.**
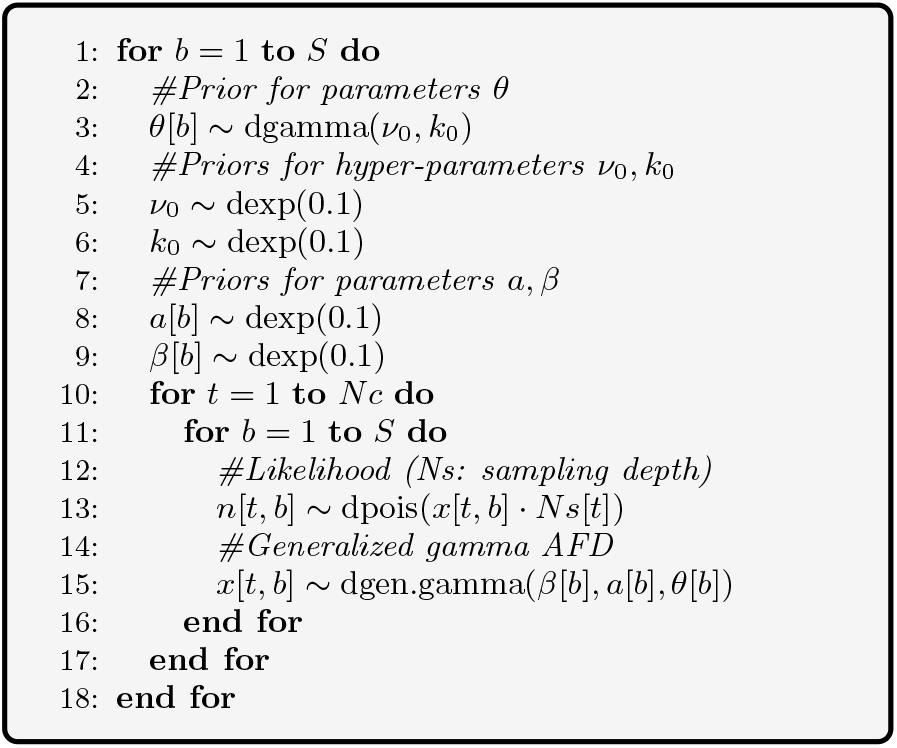
JAGS pseudocode for the hierarchical Bayesian inference described in Sec. IV.

## Appendix D: Log-fold change distribution

The log-fold change *λ* = log(*x*_*s*+1_*/x*_*s*_) is a frequently employed metric to measure these fluctuations in microbial abundances around a stationary state [17, 39]. In its definition, *x*_*s*_ and *x*_*s*+1_ denote the abundances of a taxon at consecutive samples. In our study, focused on cross-sectional data, the samples are collected from various communities: thus, *s* and *s* + 1 merely label different samples. Thus, the so-called log-fold change distribution (LFD) is a result of the AFD. In order to show this, we used the variable *u* defined by Eq. (11), in terms of which

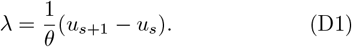

Coming from two different samples, *u*_*s*_ and *u*_*s*+1_ are independent random variables with probability density (12). Therefore,

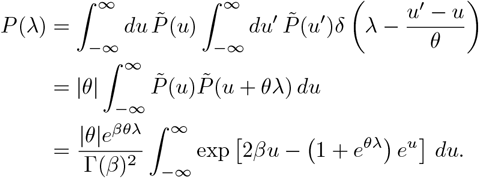

Changing to the variable *t* = (1 + *e*^*θλ*^) *e*^*u*^,

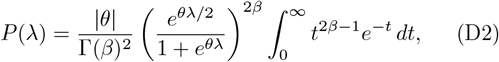

which immediately yields (27).

