## Supplementary information of the text for "Microbial populations hardly ever grow logistically and never sublinearly"

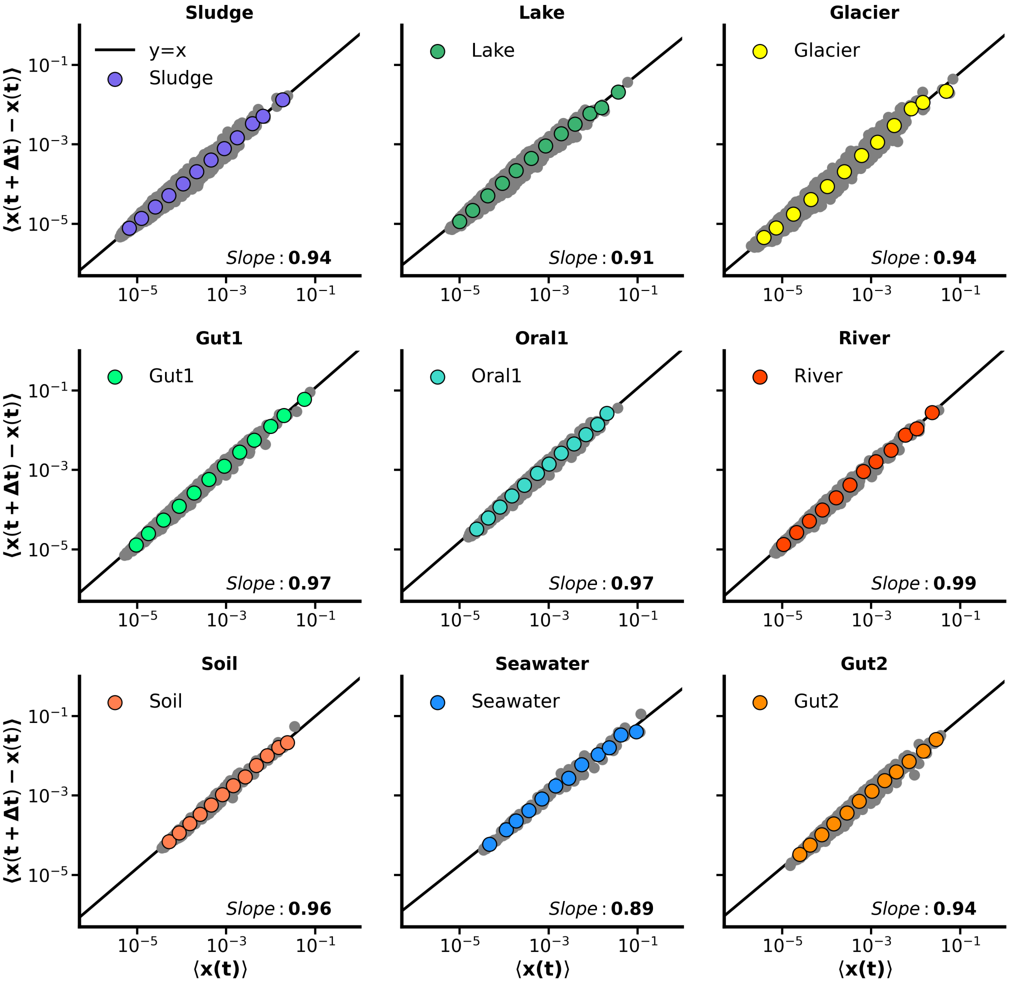


FIG. S1. **The mean absolute difference between abundances at successive time points suggests that environmental linear noise is the main source of stochasticity**. Microbial communities may be subjected by a myriad of noisy processes. In order to decide what is the main source of stochasticity we analyzed the differences between abundances at successive time points. Grey points in the background reflects the whole variability in the data sets, while the color points are a (binning) average. The figure illustrates that a model incorporating linear noise $\propto x$ most accurately reflects the experimental data.

*

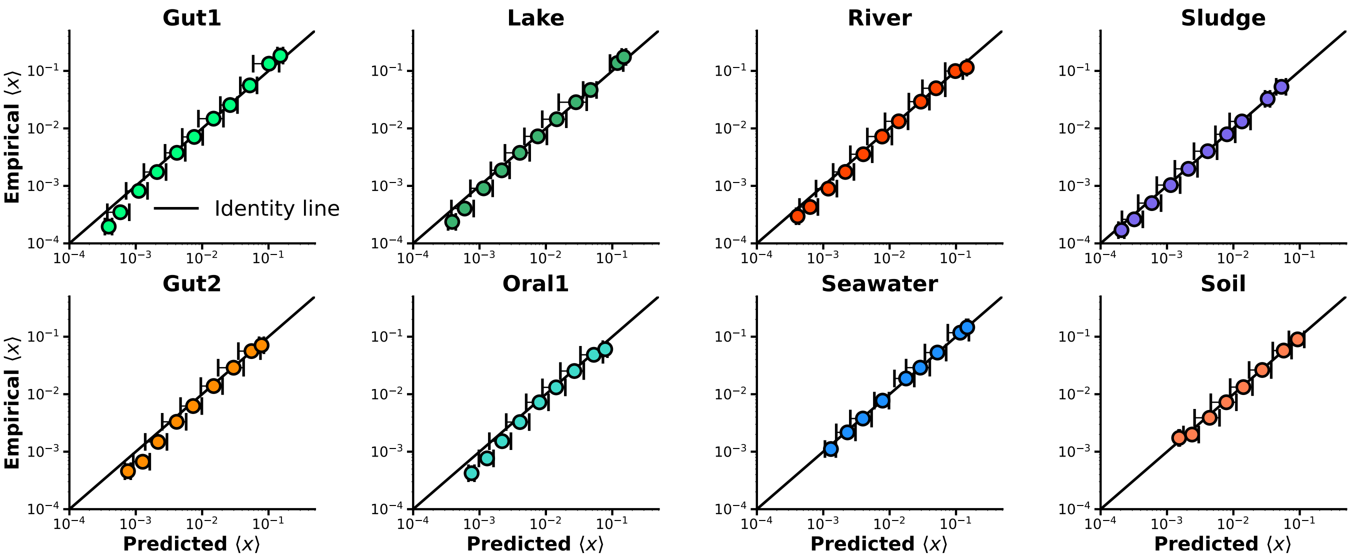


FIG. S2. **Predicted average abundance over posterior samples using the S$\theta$LM for the biome species vs. empirical ones**. Dots provide a coarse-grained visualization of the average with error bars displaying the variability within groups. For all biomes the posterior samples correctly predicts the mean abundance.

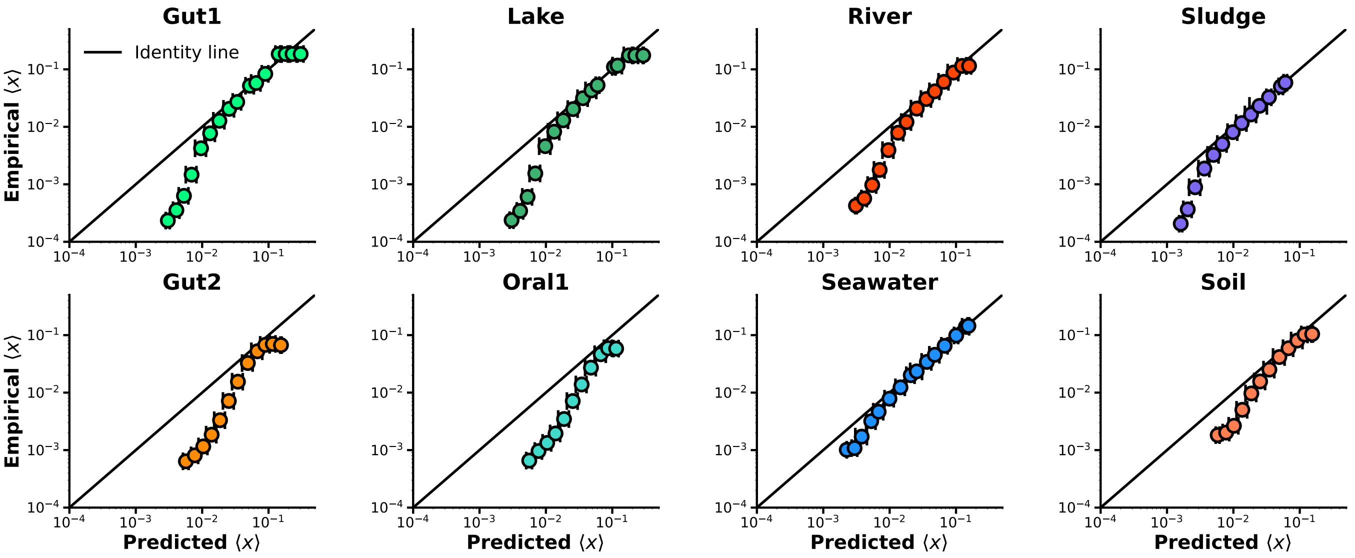


FIG. S3. **Predicted average abundance over posterior samples using the SLM for the biome species vs. empirical ones**. Dots provide a coarse-grained visualization of the average with error bars displaying the variability within groups. For all biomes the posterior samples tends to understimate the mean abundance, specially for low abundances.

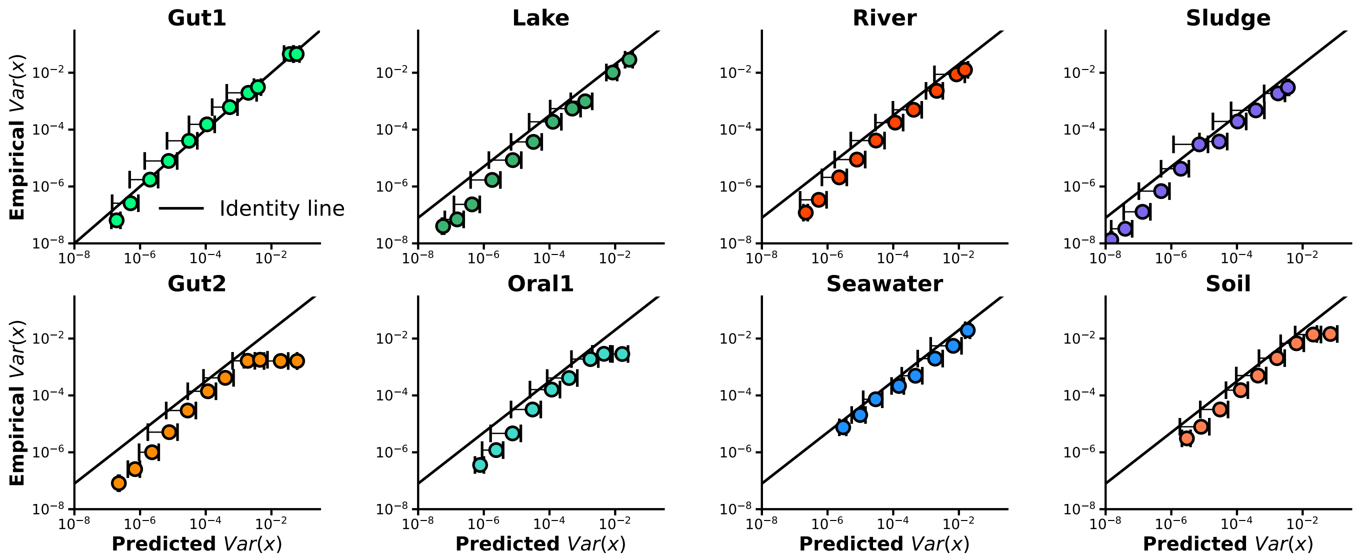


FIG. S4. **Predicted abundance variance over posterior samples using the S$\theta$LM for the biome species vs. empirical ones**. Dots provide a coarse-grained visualization of the average with error bars displaying the variability within groups. For all biomes the posterior samples correctly predicts the abundance variance.

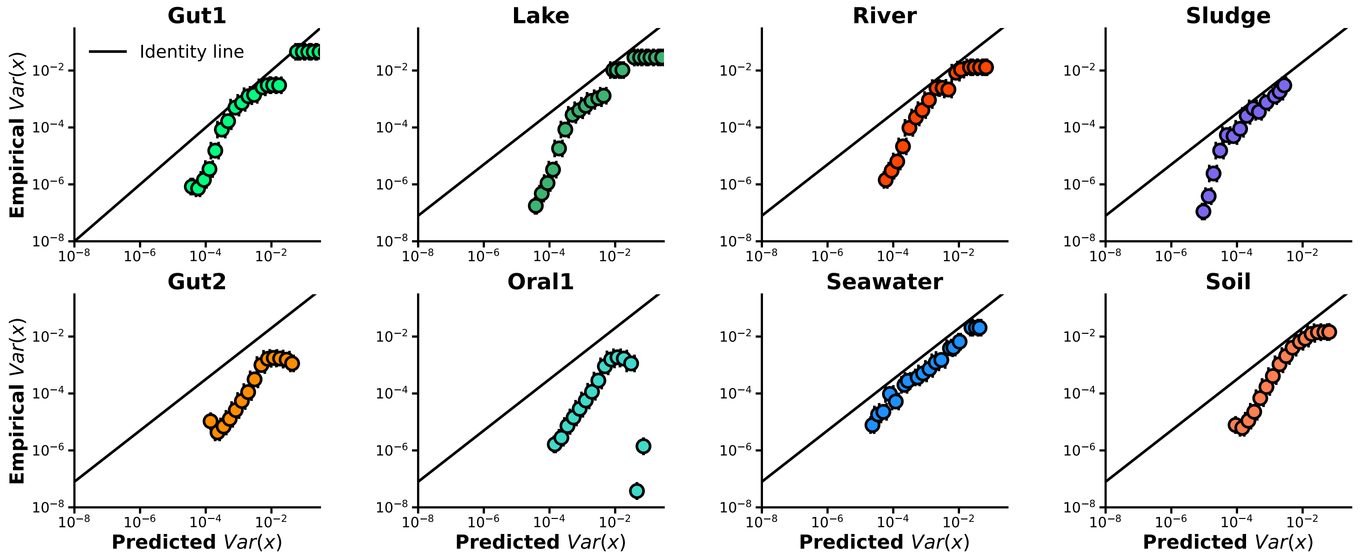


FIG. S5. **Predicted abundance variance over posterior samples using the SLM for the biome species vs. empirical ones**. Dots provide a coarse-grained visualization of the average with error bars displaying the variability within groups. For all biomes the posterior samples tends to understimate the abundance variance, specially for low abundances.

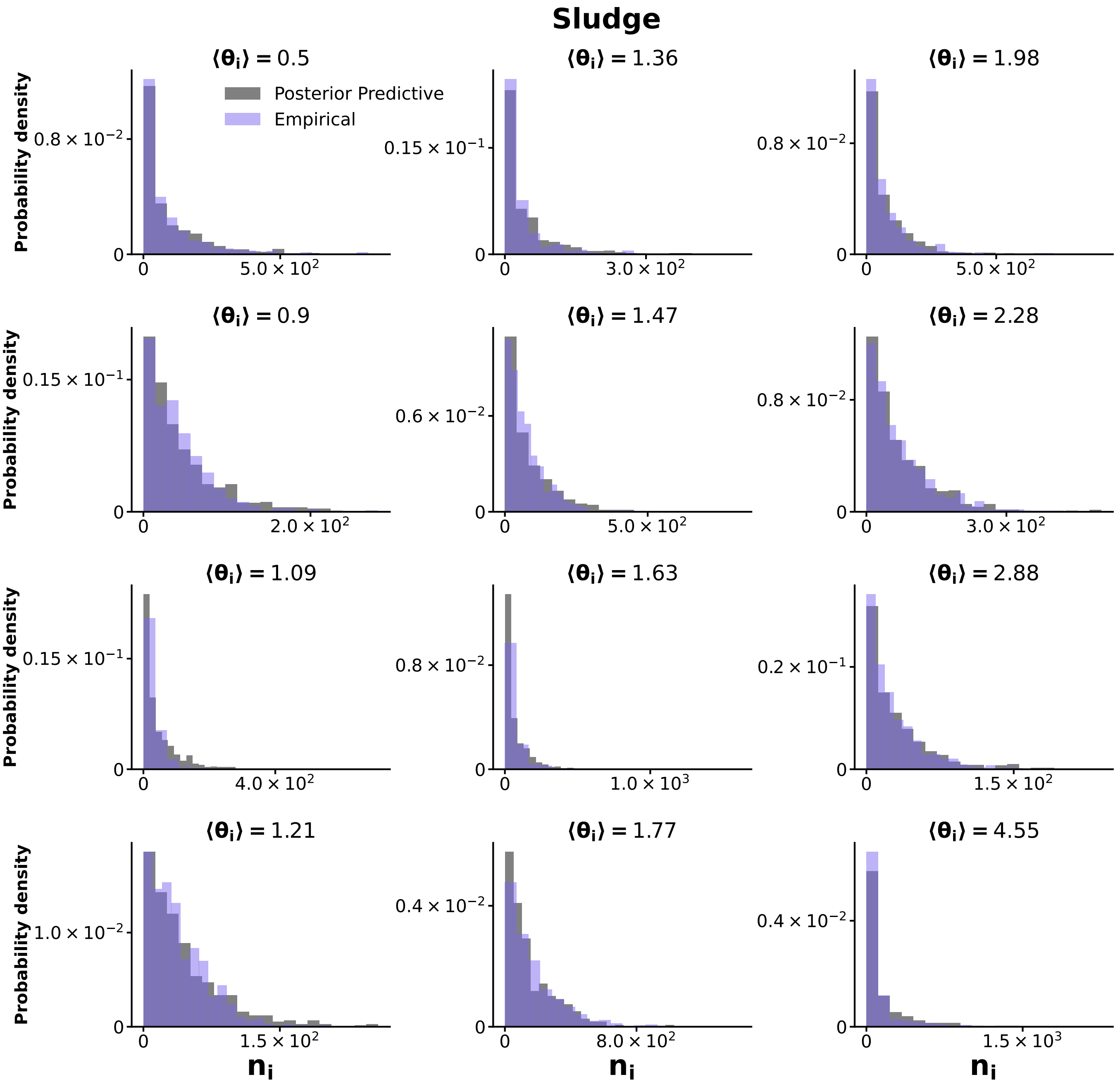


FIG. S6. **Posterior predictive tests for the Sludge biome dataset.** By comparing the posterior predictive distributions (gray) with empirical data (purple) for three different taxa we assess the accuracy of the inference. The x-axis represents the number of counts for each taxon $n_i$. We select taxons with increasing $\langle\theta_i\rangle$ to avoid any biases. The plots demonstrate the alignment between the model predictions and the observed empirical data, highlighting the accuracy of the Bayesian inference approach in modeling the abundances distributions.

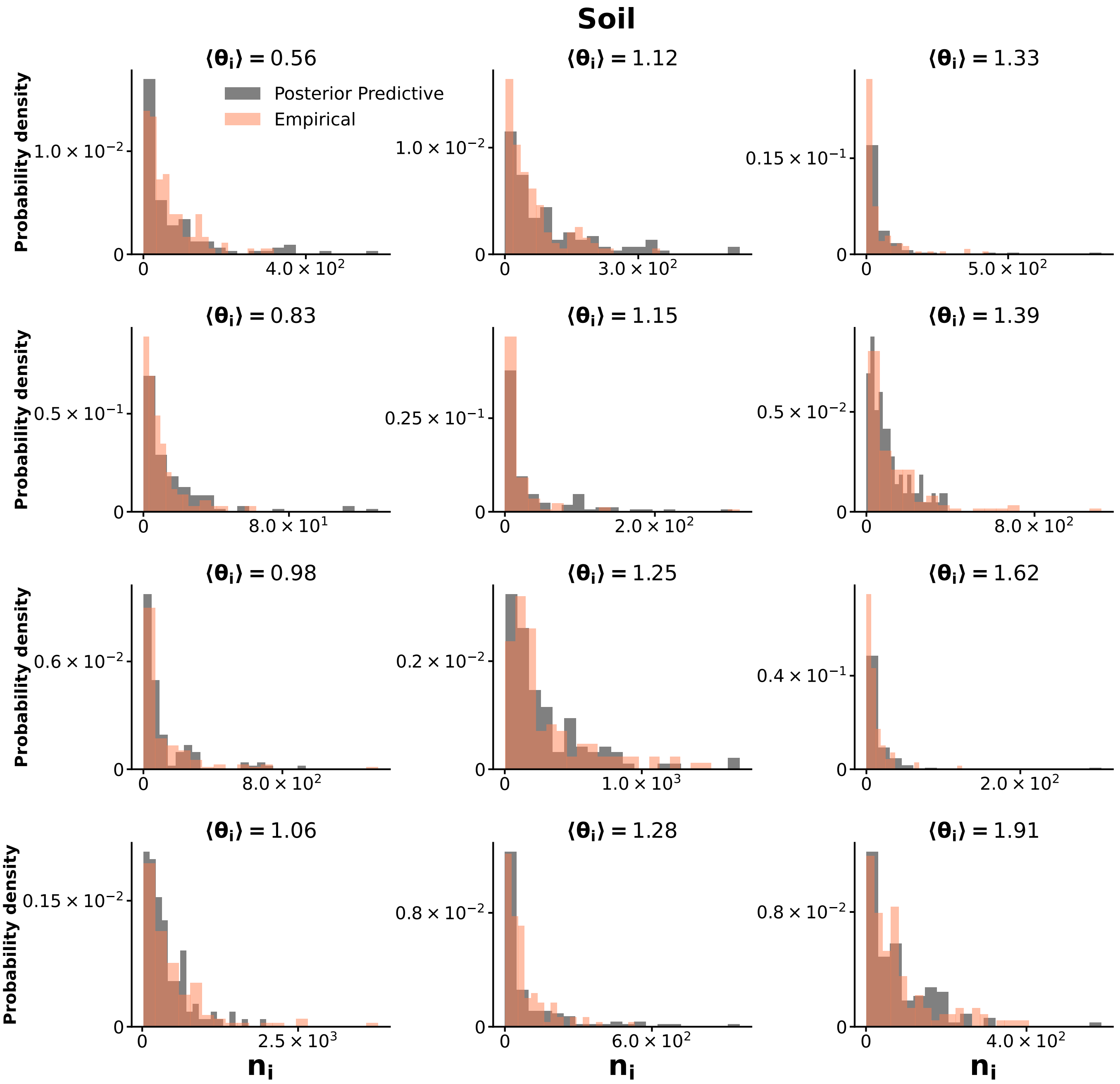


FIG. S7. **Posterior predictive tests for the Soil biome dataset.** Same as Fig. S6 using the Soil biome database (orange).

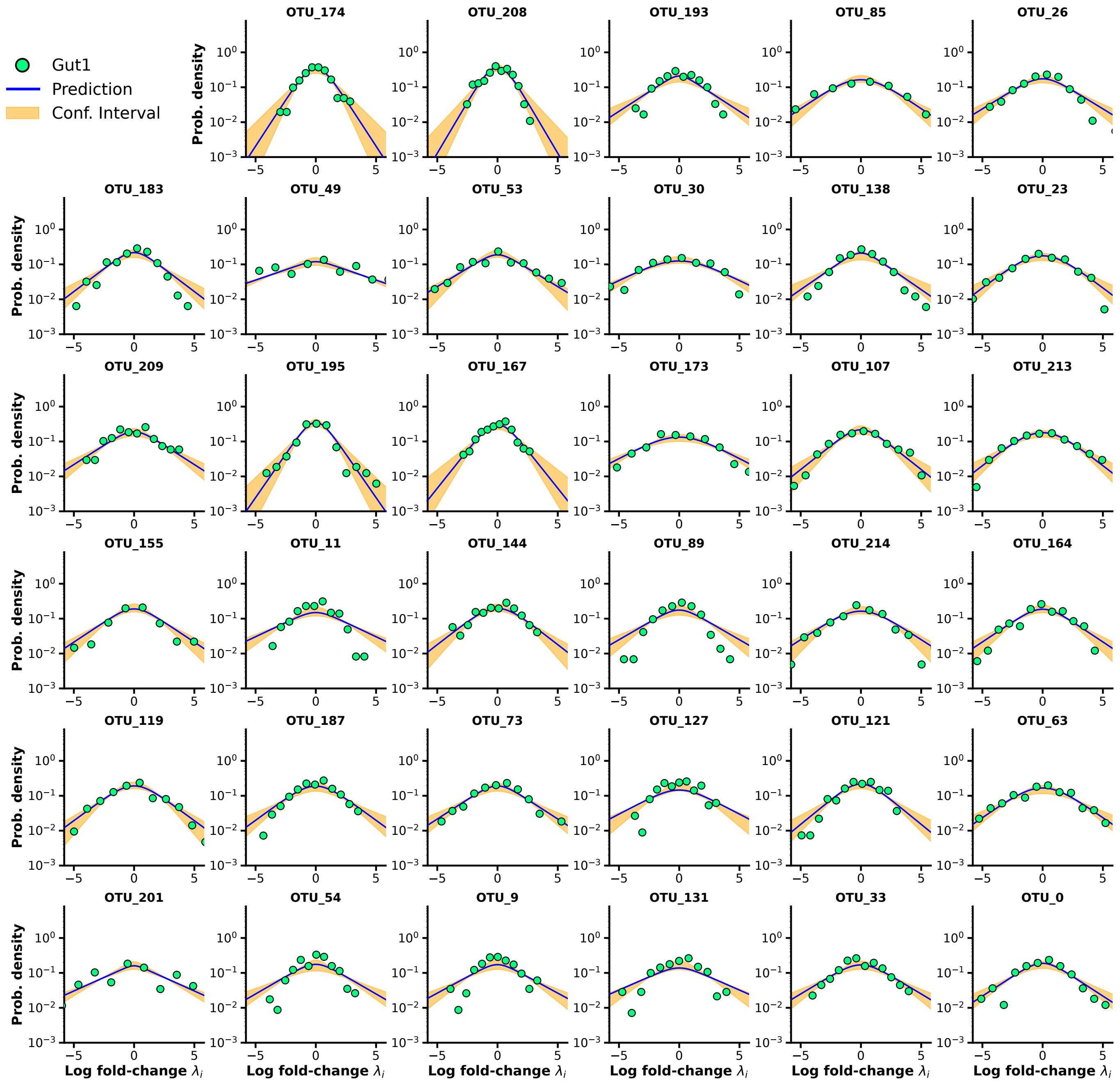


FIG. S8. **Log-fold change distribution (LFD) as a consistency test for Bayesian estimates of the Gut1 dataset**. This figure illustrates the log-fold change distribution for several randomly selected taxa, comparing empirical data (colored points) with predictions made using maximum a posteriori estimates from Eq. (27) of the main text (blue lines). The results demonstrate that the Bayesian approach accurately captures the LFD, with most empirical data points falling within the predicted confidence intervals, thereby validating the model.

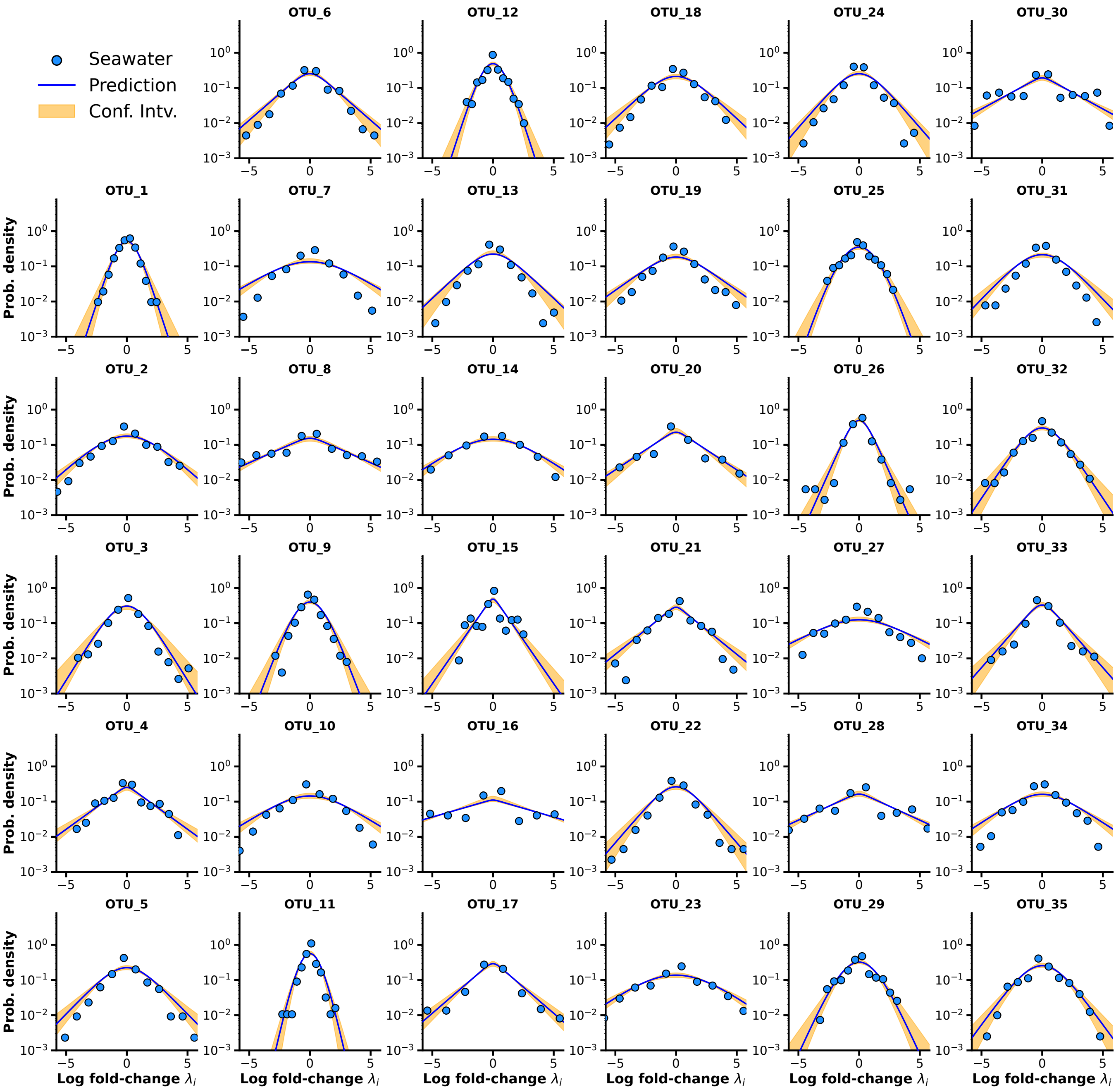


FIG. S9. **LFD as a consistency test for Bayesian estimates of the Seawater dataset**. Same as Fig. S8 using the Seawater biome database.
